# Endodermal BRD4 mediates epithelial-mesenchymal crosstalk during lung development

**DOI:** 10.1101/2023.05.21.541621

**Authors:** Derek C. Liberti, Hongbo Wen, Kwaku K. Quansah, Prashant Chandrasekaran, Josh Pankin, Nigel S. Michki, Annabelle Jin, MinQi Lu, Maureen Peers De Nieuwburgh, Lisa R. Young, Rajan Jain, David B. Frank

**Author notes:** Corresponding Author and Lead Contact: David B. Frank, M.D., Ph.D. Children’s Hospital of Philadelphia 3615 Civic Center Blvd ARC 416 K Philadelphia, PA 19104. These authors contributed equally to this work (Derek C. Liberti, Hongbo Wen).

## Abstract

Lung morphogenesis relies on diverse cell intrinsic and extrinsic mechanisms to ensure proper cellular differentiation and compartmentalization. Using genetic mouse models, tissue explants, and transcriptomic analysis, we demonstrate that the epigenetic reader Bromodomain Containing Protein 4 (BRD4) is required for lung morphogenesis and perinatal survival. Endodermal BRD4 deletion impairs epithelial-mesenchymal crosstalk, leading to disrupted proximal-distal patterning and branching morphogenesis. These early defects result in dilated airways and the formation of cystic distal airway structures, containing both airway and alveolar features. Moreover, BRD4 deficient lungs exhibit abnormal airway and alveolar epithelial cell lineage allocation and differentiation. Restoration of SHH signaling partially rescues defects due to loss of BRD4, suggesting a role for BRD4 in regulating the SHH-FGF signaling axis. Together, these data identify the essential role of Brd4 in lung development to ensure proper intercellular crosstalk to enable proper lineage specification, identity, and maturation.

## INTRODUCTION

Lung development relies on precise, highly stereotyped intercellular communication networks to establish the complex architecture of the adult organ. In particular, epithelial-mesenchymal signaling guides branching morphogenesis via well investigated molecular pathways, including SHH, FGF, BMP, retinoic acid, TGF-β, and WNT.^1–4^ Lung morphogenesis additionally relies on epigenetic factors, such as DNA methyltransferases, histone modification enzymes, and noncoding RNAs, to ensure the proper epithelial lineage specification, proliferation, and maturation necessary to generate a functional lung.^5–14^ Epithelial cell fate decisions occur in a spatially specific manner and are major drivers of not only lung development but also adult regeneration after injury.^15–17^ The ability of epigenetic factors to manipulate epithelial cell fate has made them attractive targets for amelioration of human disease, but our understanding of these factors remains limited.^18, 19^ Elucidating the role of epigenetic factors in epithelial cell fate is essential to develop targeted therapies to improve lung disease outcomes.

Bromodomain containing protein 4 (BRD4) is an epigenetic reader with multiple roles in transcription regulation. BRD4 has been shown to serve as a scaffold to recruit transcription factors to acetylated histones, act as a kinase and histone acetyltransferase, and maintain chromatin architecture.^20–24^ Due to its broad functionality, BRD4 is essential for early embryonic mouse development and pluripotent stem cell identity maintenance.^25, 26^ In addition, BRD4 is a key regulator of normal cell cycle progression.^27, 28^ BRD4 has been implicated in the potentiation of pulmonary arterial hypertension and pulmonary fibrosis, but the role of this factor during lung development is largely unknown.^29–31^

Using *in vivo* genetic mouse models, *ex vivo* lung explant assays, and single-cell transcriptomics, we investigated the role of BRD4 in the developing lung endoderm. We show that endodermal deletion of BRD4 results in the emergence of cystic distal airway structures and perinatal lethality due to respiratory failure. Further, we demonstrate that BRD4 deficient mice exhibit impaired lung branching, resulting in fewer, dilated proximal airways and enlarged distal airspaces. Perturbed morphogenesis appears to result from a loss of epithelial SHH and increased and expanded mesenchymal FGF10 signaling at the distal branching tips. Mutant mice display a loss of airway and alveolar epithelial cell maturation, particularly within cystic distal airway structures, which are encased in ectopic smooth muscle. Ultimately, we show that activation of SHH is sufficient to partially rescue the cystic distal airway phenotype. Taken together, our data demonstrate that BRD4 acts upstream of SHH to promote proper epithelial lineage determination and localization.

## RESULTS

### Endodermal BRD4 is required for lung morphogenesis and perinatal survival

To determine the role of BRD4 in the developing lung, we generated *Shh^Cre^;Brd4^flox/flox^* mice, allowing conditional deletion of BRD4 in the lung endoderm, and compared these mutant mouse lungs to controls at E12.5, E15.5, and E18.5 (**Figure 1A**).^23, 32^ While we detected BRD4 protein throughout the developing embryo, we confirmed complete loss of BRD4 from the lung endoderm in *Shh^Cre^;Brd4^flox/flox^* mice at E12.5 (**Figure 1B**). BRD4 deficient lungs exhibited abnormal morphogenesis, including the formation of cystic distal airway structures (**Figure 1C**). These cystic distal airway structures were apparent in mutant lungs at E15.5, along with dilated airways, which were more pronounced at E18.5 (**Figure 1D**). Additionally, morphological analysis of tissue sections at E18.5 revealed statistically significant expansion of alveolar airspaces in mutant lungs compared to controls (**Figures 1D-F**). *Shh^Cre^;Brd4^flox/flox^* mice did not survive until weaning, and we found that *Shh^Cre^;Brd4^flox/flox^* mice became cyanotic and perished shortly after birth, likely due to respiratory distress (**Figures S1A and S1B**). While mutant lungs floated, suggesting *Shh^Cre^;Brd4^flox/flox^*mice could inhale, these lungs floated due to collection of large air bubbles in their cystic airway structures (**Figure S1C**). Together, these data show that endodermal BRD4 is required for lung development.

**Figure 1:**
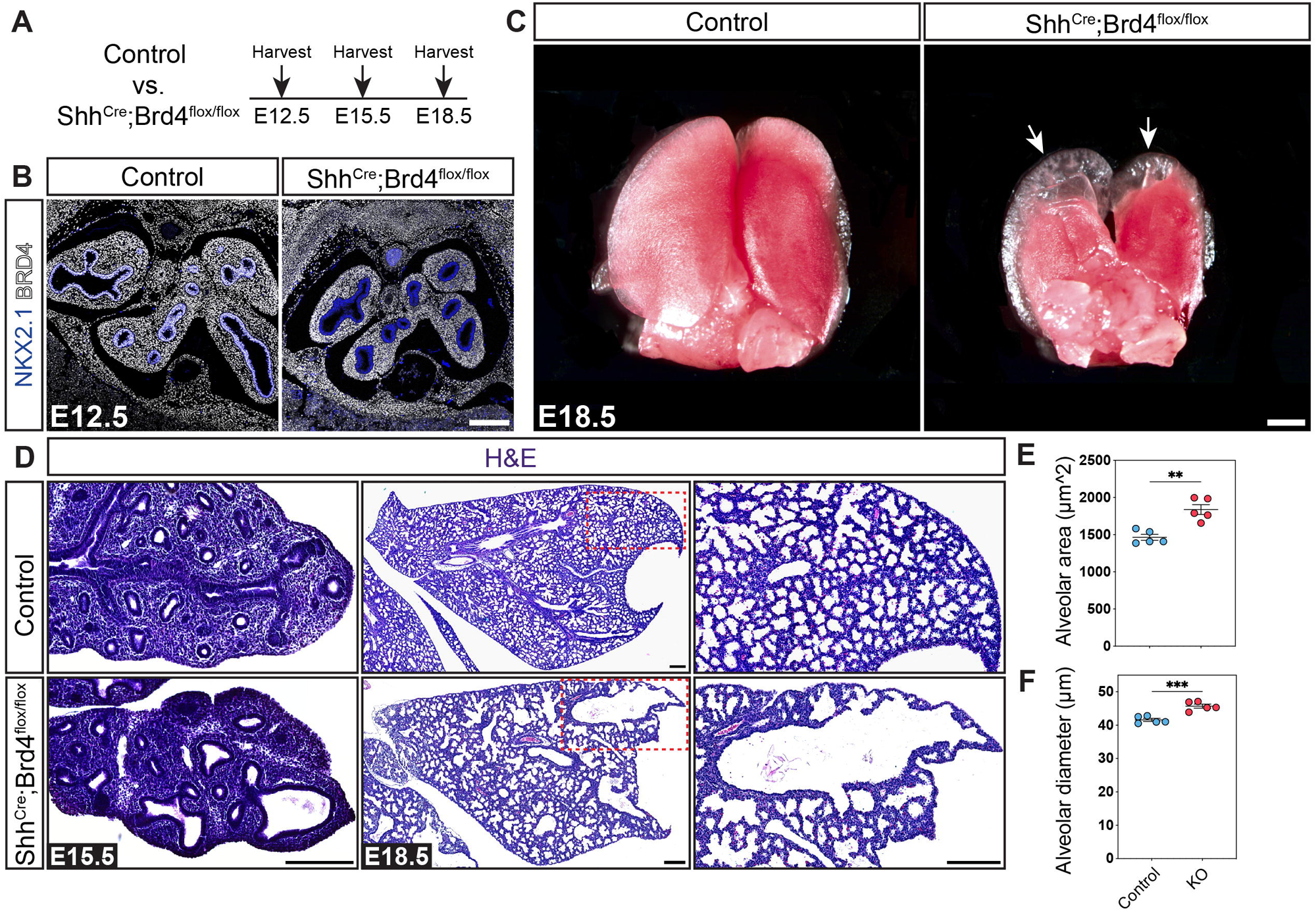
Endodermal BRD4 is required for lung morphogenesis and perinatal survival. (A) Experimental schematic indicating the timepoints analyzed for data shown. (B) Immunohistochemistry (IHC) for NKX2.1 and BRD4 at E12.5 (scale bar: 200μm). (C) Brightfield images of whole control and mutant lungs at E18.5. White arrows mark cystic distal airway structures (scale bar: 2mm). (D) Hematoxylin and eosin (H&E) staining at E15.5 and E18.5 (all scale bars: 200μm). (E-F) Quantification of morphological changes shown in panel D at E18.5. Data are represented as mean ± SEM. Two-tailed t test: **p ≤ 0.01, ***p ≤ 0.001, n = 5.

### Loss of endodermal BRD4 disrupts epithelial-mesenchymal crosstalk

To assess how loss of BRD4 results in the formation of cystic distal airway structures, we investigated early branching morphogenesis in control and mutant lungs. As early as E12.5, we observed impaired branching in BRD4 deficient mice (**Figures 2A-B**). To determine whether the observed defect was lung intrinsic, we harvested control and mutant lungs at E12.5 and cultured them *ex vivo* for three days. Mutant lungs exhibited fewer and dilated airways with diminished SOX2 expression in distal airways in addition to the formation of cystic distal airway structures, consistent with our *in vivo* observations (**Figures 2C-D**). However, proximal airway dilation was more dramatic *in vivo* (**Figure 2D**). We did not observe any obvious differences in proliferation of proximal or distal epithelial cells or cell death in control or mutant mice (**Figures 2D-E and S2A)**. These data suggest that altered mutant lung morphogenesis is due to improper proximal-distal patterning.

**Figure 2:**
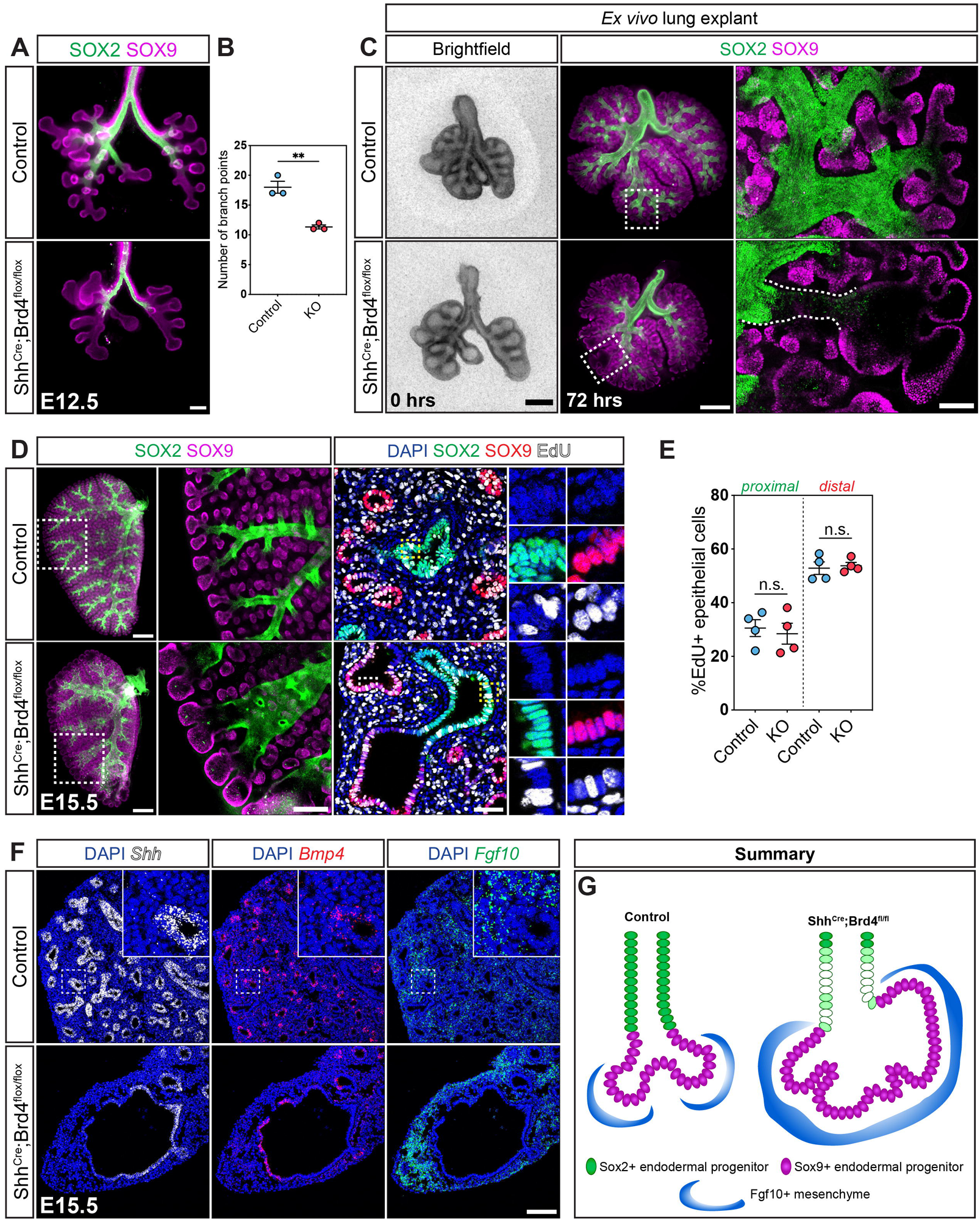
Loss of endodermal BRD4 disrupts epithelial-mesenchymal crosstalk. (A-B) Wholemount immunostaining for SOX2 and SOX9 to assess branching at E12.5 (scale bar: 200μm). Quantification data are represented as mean ± SEM. Two-tailed t test: **p ≤ 0.01, n = 3. (C) Left: brightfield images of control and mutant lungs at E12.5 (scale bar: 500μm). Middle: wholemount immunostaining for SOX2 and SOX9 after 3 days of culture ex vivo (scale bar: 500μm). Dashed white boxes indicate branching tips in control and mutant lungs. Right: zoomed in images of branching tips, dashed white lines outline a distal airway (scale bar: 100μm). (D) Left: wholemount immunostaining for SOX2 and SOX9 at E15.5 (scale bar: 500μm). Dashed white boxes marked zoomed in areas shown in middle images (scale bar: 100μm). Right: IHC for SOX2 and SOX9 and staining for EdU at E15.5. Dashed boxes mark zoomed in areas of proximal (yellow) and distal (white) endoderm (scale bar: 50μm). (E) Quantification and comparison of proximal (SOX2+EdU+/SOX2+) and distal (SOX9+EdU+/SOX9+) endodermal EdU incorporation at E15.5 in control versus mutant lungs as shown in panel (D, right). Quantification data are represented as mean ± SEM. Two-tailed t tests: ns: not significant, n = 4. (F) RNA FISH for *Shh, Bmp4*, and *Fgf10* in control and mutant E15.5 lungs (scale bar: 100μm). (G) Summary: In BRD4 mutant distal tips, downregulation of *Shh* expression and expansion of *Fgf10* is associated with a reduction in SOX2 and disruption of proximal-distal patterning and branching.

To determine whether epithelial-mesenchymal communication is altered in BRD4 deficient lungs, we investigated the expression of critical branching factors *Shh*, *Bmp4*, and *Fgf10* using multiplexed RNA fluorescent in situ hybridization (FISH) (**Figure 2F**). The SHH-BMP4-FGF10 signaling axis has been well-characterized in the lung with feedback loops between SHH and FGF signaling.^33–37^ *Shh* is typically expressed throughout the developing lung endoderm (**Figure 2F**). However, in BRD4 deficient mice, *Shh* expression was diminished at the distal tips of cystic distal airway structures (**Figure 2F**). Moreover, while *Bmp4* expression was restricted to the distal tips of airway structures in control lungs, *Bmp4* expression dramatically expanded in BRD4 deficient mice, demonstrating an enlargement of the distal tip region (**Figure 2F**). Additionally, *Fgf10* expression was increased and expanded in BRD4 mutant lungs (**Figure 2F**). Similar findings were observed in a recent study in lung development revealing that FGF singling components including ETV5 mediate the FGF-SHH signaling loop.^34^ Thus, we performed FISH for *Etv5* and *Fgfr2* but observed no changes in epithelial expression between controls and mutants (**Figures S2B**). Together, our data show that loss of BRD4 leads to impaired epithelial-mesenchymal crosstalk resulting in impaired branching morphogenesis, dilated airways, and cystic expansion of distal branching tips (**Figure 2G**).

### SHH activation partially rescues BRD4 mutant cystic distal airway structure phenotype

To further characterize the potential mechanism of SHH and FGF signaling in BRD4 mutants, we activated SHH signaling with Purmorphamine (PMA) or inhibited FGFR signaling using Alofanib, a small molecule inhibitor of FGFR, in our *ex vivo* lung explants.^38^ Treatment of mutant explants with PMA partially rescued the phenotype with a reduction in the cystic distal airway structures and expansion of SOX2 expression (**Figure 3A-B)**. However, this partial rescue was accompanied by a reduction in the number of branching tips and an increase in mesenchymal area (Figures 3D-E). Quantification of area of SOX2 expression revealed statistically significant increase with the re-emergence of SOX2 expressing airway cells in the mutant lung with PMA treatment (**Figure 3C**). Further morphometric analysis of tip number showed significant reduction in both control and mutant group with PMA treatment (**Figure 3D**). There was a trend that PMA treatment reduced tip thickness for the BRD4 deficient explant, but this did not reach statistical significance due to limited number of biological replicate (**Figure 3E**). On the other hand, treatment with Alofanib impaired branching morphogenesis in both the control and BRD4 mutant explants, suggesting intact FGF signaling in BRD4 mutants (**Figures S3A-B**). Thus, BRD4 regulates lung morphogenesis and proximal-distal patterning at the bronchoalveolar junction, in part, via regulation of SHH signaling (**Figure 3F**).

**Figure 3:**
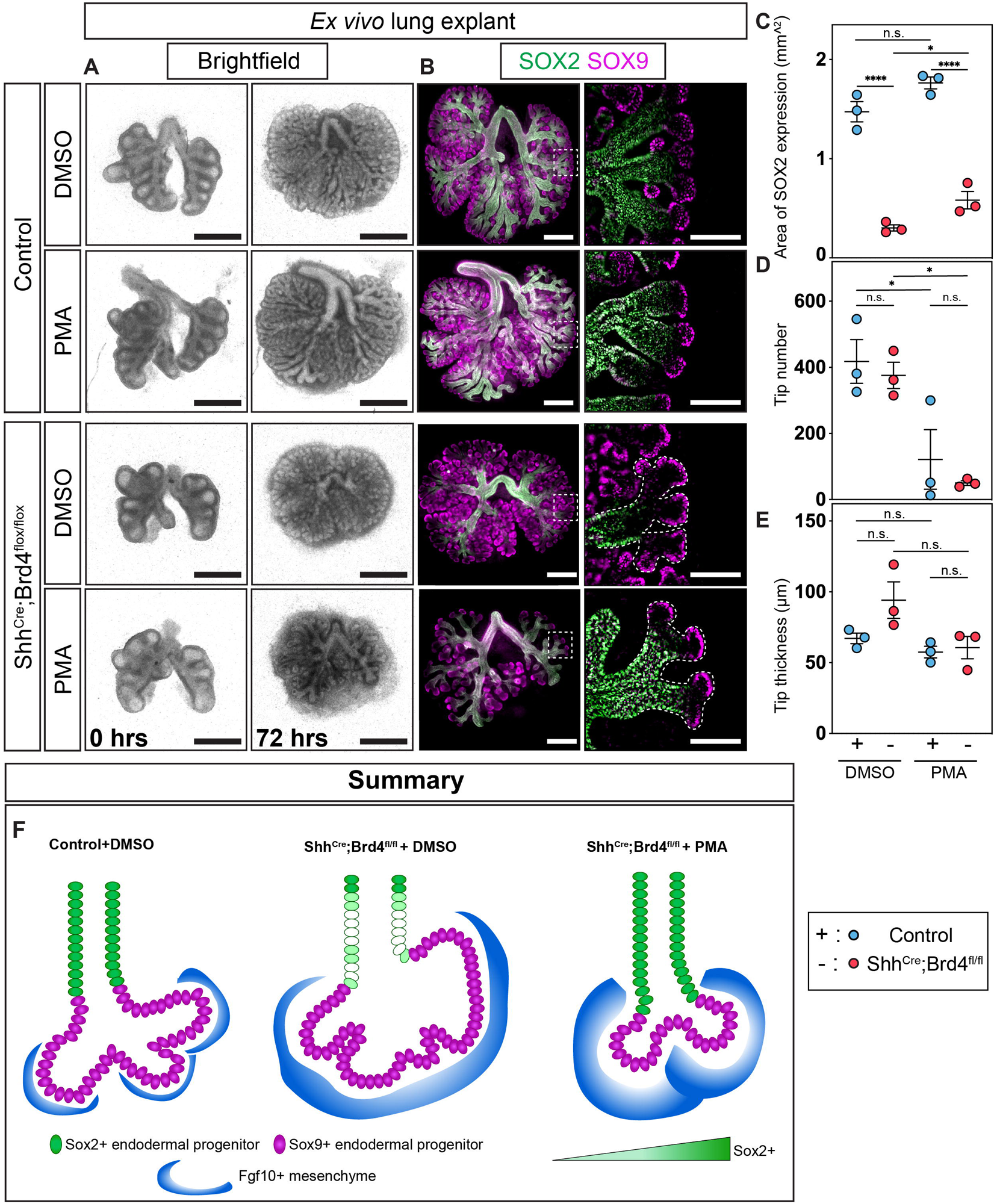
Shh activation partially rescues BRD4 mutant cystic distal airway structure phenotype. (A) *Ex vivo* lung explant brightfield images of control and mutant lungs at (left) E12.5 and (right) E15.5 with treatments: DMSO and Purmorphamine (PMA) (scale bar: 500μm). (B) Left: wholemount immunostaining for SOX2 and SOX9 after 3 days of *ex vivo* culture (scale bar: 500μm). Dashed white boxes indicate branching tips in control and mutant lungs. Right: zoomed in images of branching tips, dashed white lines outline a distal airway (scale bar: 250μm). (C-E) Quantification and comparison of (C) area of SOX2 expression, (D) tip number and (E) tip thickness between control and mutant lungs treated with DMSO and PMA. Quantification data are represented as mean ± SEM. Two tailed t-tests: n.s.: not significant, *p ≤ 0.05, ****p ≤ 0.001, n = 3. (F) Summary: Treatment with the sonic hedgehog pathway activator, PMA, partially rescues aberrant development in BRD4 mutant lungs. There is expansion of SOX2 and a reduction in the size of the distal cystic structures.

### Endodermal BRD4 is essential for epithelial lineage segregation and differentiation

To determine how early patterning defects in BRD4 mutants impact lung maturation, we examined the composition of the cystic distal airway structures. The apical surfaces of these structures were lined with cuboidal epithelial cells (**Figure 4A**). Yet, these structures lacked a distinct bronchoalveolar duct junction, leading to poor lineage allocation and maturation of airway and alveolar epithelial cells (**Figures 4A-B**). Epithelial cells in the proximal portion of cystic distal airway structures weakly expressed markers for secretory and multiciliated cells, including SCGB1A1 and TUBB4, while cuboidal epithelial cells in the distal portion expressed alveolar epithelial type 1 (AT1) and type 2 (AT2) markers, including HOPX and SFTPC (**Figures 4A-B**). However, HOPX expression expanded into the proximal region of the cystic distal airway structures, and some SFTPC+ cells also expressed HOPX (**Figures 4A-B**). Interestingly, we also observed ectopic basal or basaloid-like cells within the alveolar region in BRD4 mutant mice (**Figure S4A**). In addition to improper epithelial lineage allocation and maturation, these cystic distal airway structures were encased in ectopic smooth muscle, suggesting impaired epithelial-mesenchymal crosstalk (**Figures 4C-D**).

**Figure 4:**
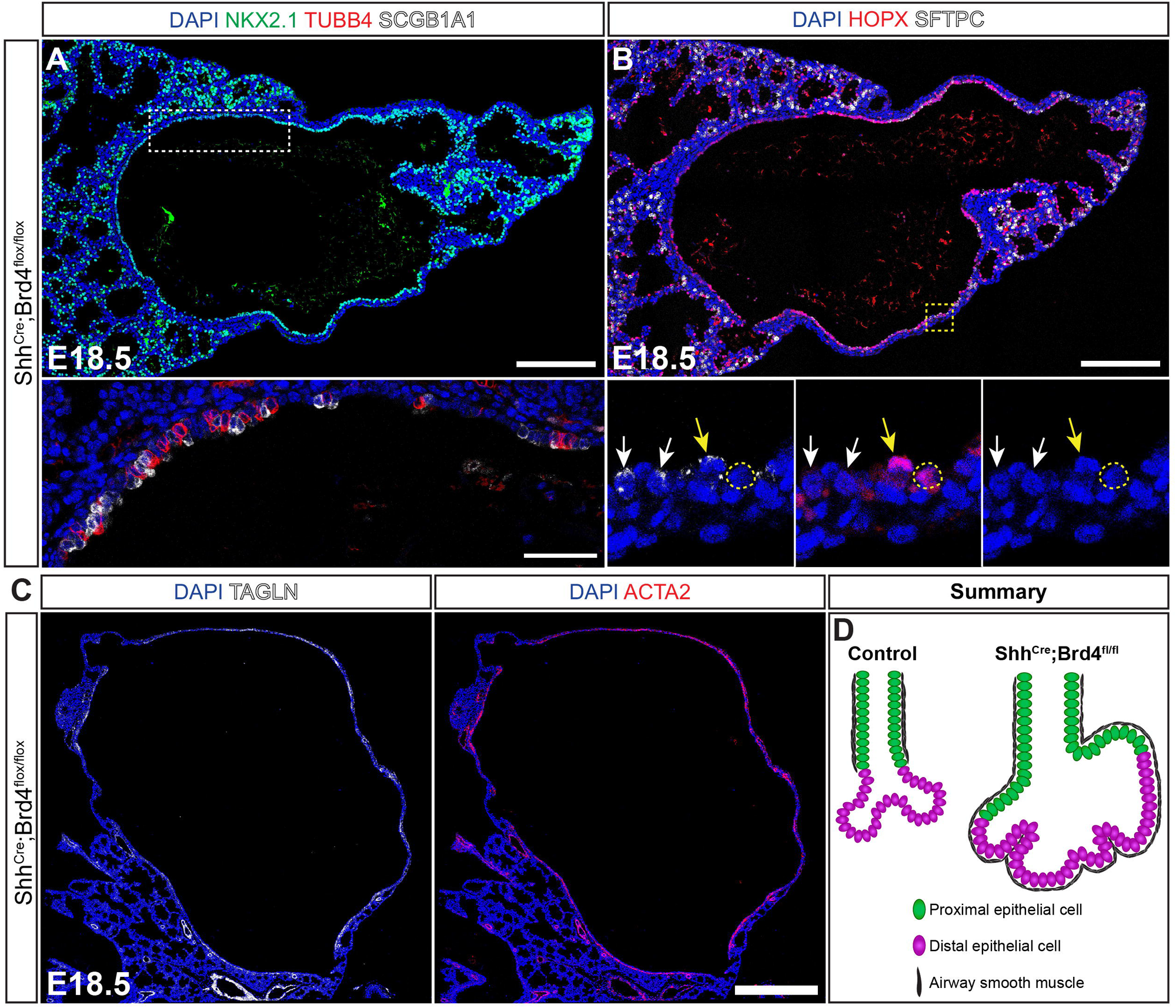
Endodermal BRD4 is essential for epithelial lineage segregation and differentiation. (A) IHC for NKX2.1, TUBB4, and SCGB1A1 in BRD4 mutant E18.5 lungs (scale bar: 200μm). Dashed white box marks the zoomed in region shown in the lower image (scale bar: 50μm). (B) IHC for HOPX and SFTPC (scale bar: 200μm). Dashed yellow box marks the zoomed in region shown in the lower image. Dashed yellow circles mark AT1 cells, white arrows mark AT2 cells, and yellow arrows mark HOPX+ SFTPC+ cells. (C) IHC for TAGLN and ACTA2 in Brd4 mutant lung cystic distal airway structures at E18.5. (D) Summary: Proximal and distal epithelial cells become poorly segregated in cystic distal airway structures with loss of the bronchoalveolar duct junction, impaired epithelial differentiation, and ectopic smooth muscle.

### BRD4 is required for secretory cell differentiation and maintenance of airway epithelial cell identity at the bronchoalveolar duct junction

Although BRD4 mutant airways often transitioned abruptly into cystic distal airway structures, some appeared to terminate into relatively normal bronchoalveolar duct junctions, allowing for continued formation of the alveolar compartment (**Figure 5A**). Although secretory cells continued to express SCGB1A1 in the proximal intralobular airways, they expressed markedly less SCGB1A1 protein than controls in both proximal and distal regions, particularly within cystic distal airway structures (**Figures 5A-B**). Similarly, both proximal and distal multiciliated cells appeared to lose TUBB4 marker gene expression (**Figures 5A-B**). We performed quantification of the number of secretory and multiciliated cells in both proximal and distal compartments. There was approximately a 50% percent reduction in the number of secretory cells in both proximal and distal regions (**Figure 5C**). Despite maintaining marker expression in proximal airways, the number of multiciliated cells in BRD4 mutants decreased by about 40% percent proximally and 50% distally compared to controls (**Figure 5D**).

**Figure 5:**
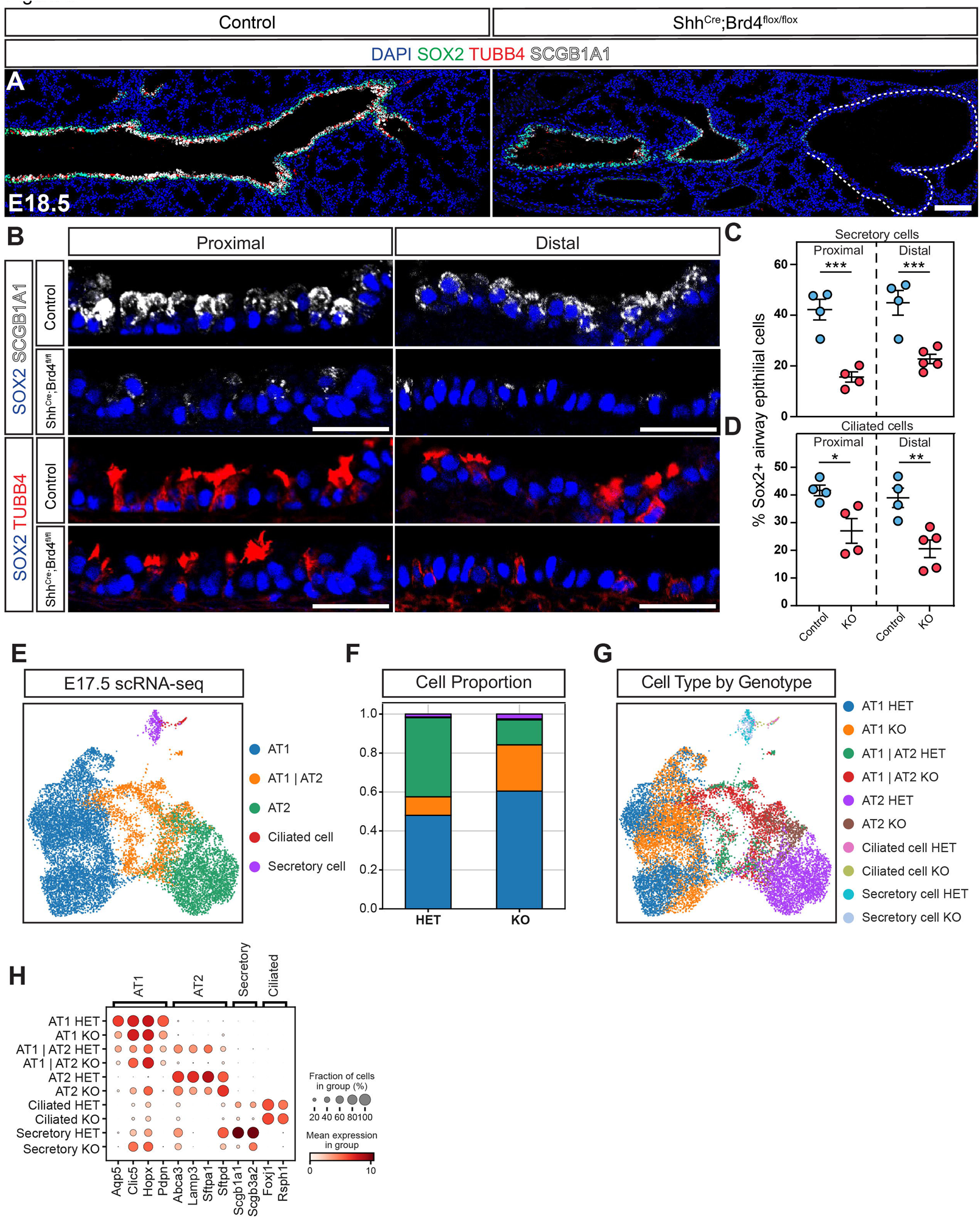
BRD4 is required for secretory cell differentiation and maintenance of airway epithelial cell identity at the bronchoalveolar duct junction. (A) IHC for SOX2, TUBB4, and SCGB1A1 control and mutant E18.5 lungs (scale bar: 100μm). Dashed white line outlines a cystic distal airway structure. (B) IHC for SOX2, TUBB4, and SCGB1A1 in proximal and distal intralobular airway regions of control and BRD4 mutant E18.5 lungs. For BRD4 mutants, distal images reflect airways terminating into cystic distal airway structures (scale bar: 25μm). (C-D) Quantification of number of secretory (C) and ciliated (D) cells in the proximal and distal intralobular airway regions. Quantification data are represented as mean ± SEM. Two tailed t-tests: *p ≤ 0.05, **p ≤ 0.01, ***p ≤ 0.005, ****p ≤ 0.001, n ≥ 4. (E) UMAP embedding of epithelial cell clusters in BRD4 heterozygous lungs. (F) Cell compositional changes in BRD4 heterozygous and mutant lungs based on cell proportion. (G) UMAP embedding of epithelial cell clusters separated by cell type in BRD4 heterozygous lungs and mutant lungs. (H) Dot plot of marker genes for each cell type where dot size indicates the proportion of cells within a cluster expressing a gene, and color intensity indicates the relative expression level.

To explore how loss of BRD4 impacts epithelial cell differentiation globally, we isolated heterozygous control and homozygous BRD4 deficient epithelial cells at E17.5 and performed single-cell RNA sequencing (scRNA-seq) to compare transcriptional and cellular changes (**Figure 5E**). Using fluorescence activated cell sorting (FACS), we collected EPCAM+ cells from combined heterozygous control or homozygous embryos in each litter and sequenced using the 10X Genomics Chromium droplet-based scRNA-seq platform. After quality control, we sequenced over 6,520 control and 7,151 homozygous knockout cells. Cell clustering and cell annotation confirmed the presence of major intrapulmonary epithelial cell populations in the lung, including AT1, AT2, secretory, and multiciliated airway cells. Total cell number and cell proportion of epithelium differed between control and BRD4 mutant mice (**Figures 5F and S5A**). In BRD4 mutants, there was a loss of AT2s concomitant with a gain in AT1s and in cells simultaneously expressing markers of AT1s and AT2s (AT1/A2 cells), suggesting loss of cell identity. UMAP embedding by genotype and cell type showed predominantly segregated cell clustering, indicating significant transcriptome changes with loss of BRD4 (**Figure 5G**). In addition, profiling marker gene expression in control and mutant samples revealed a loss of mature marker gene expression across AT1, AT2, secretory, and multiciliated cells (**Figure 5H**). Thus, our data demonstrate that BRD4 is required for maintenance of the bronchoalveolar duct necessary for proximal-distal lineage allocation and maturation.

Given the effects of BRD4 loss on secretory and multiciliated cells, we examined the transcriptomes of each cell type and investigated gene expression and function. Assessment of top marker genes for secretory cells revealed a significant decrease in *Scgb1a1* and *Scgb3a2* expression (**Figure S5B**). Over-representation analysis (ORA) of the transcriptome of secretory cells from BRD4 mutant versus control mice revealed enrichment in genes for protein synthesis but deficiencies in regulation of lipid metabolic processes and intracellular transport (**Figures S5C-D**). Together, ORA analysis and gene expression suggested a possible functional deficit in secretoglobin secretion. While ORA assessment of the transcriptome in multiciliated cells in BRD4 mutants identified enrichment of genes for cell death and cell cycle regulation, our quantification of proliferation and apoptosis did not demonstrate these global changes (**Figures S5E and 2E**). On the other hand, there was a decrease in enrichment of genes important for protein polymerization, musculoskeletal movement, fiber organization, and cytoskeletal organization, processes that have been described important in ciliogenesis (**Figure S5F**).^39^ We also assessed multiple signaling pathways and stress responses using Hallmark scoring on gene set enrichment analysis and determined that secretory cells downregulated many signaling pathways and were enriched in genes for stress responses (**Figure S5H**). These findings demonstrate significant transcriptomic and cell number changes affecting secretory cells, indicating a global functional deficit in them.

### BRD4 is critical for AT2 lineage determination and identity and AT1 cell maturation

The cell proportion analysis from our scRNA-seq data suggested significant defects in cell identity and specification, with a decrease in AT2 and increase in AT1/AT2 cells (**Figures 5E-G and S5A**). To assess how loss of BRD4 impacts alveolar epithelial cell differentiation, we investigated gene expression differences between control and mutant AT1 and AT2 cells using our scRNA-seq data. ORA of AT1s identified downregulation of genes associated with cell extension and contraction and upregulation of genes modulating cellular metabolism (**Figures 6A and S6A**). ORA of control versus BRD4 deficient AT2 cells revealed alterations in genes involved in lipid metabolism, synthesis, and transport (**Figure 6B**). Interestingly, AT2s became more enriched in AT1 marker genes and pathways involved in vascular development, indicating a potential transition into an AT1 state (**Figures S6B and 6C-D**).

**Figure 6:**
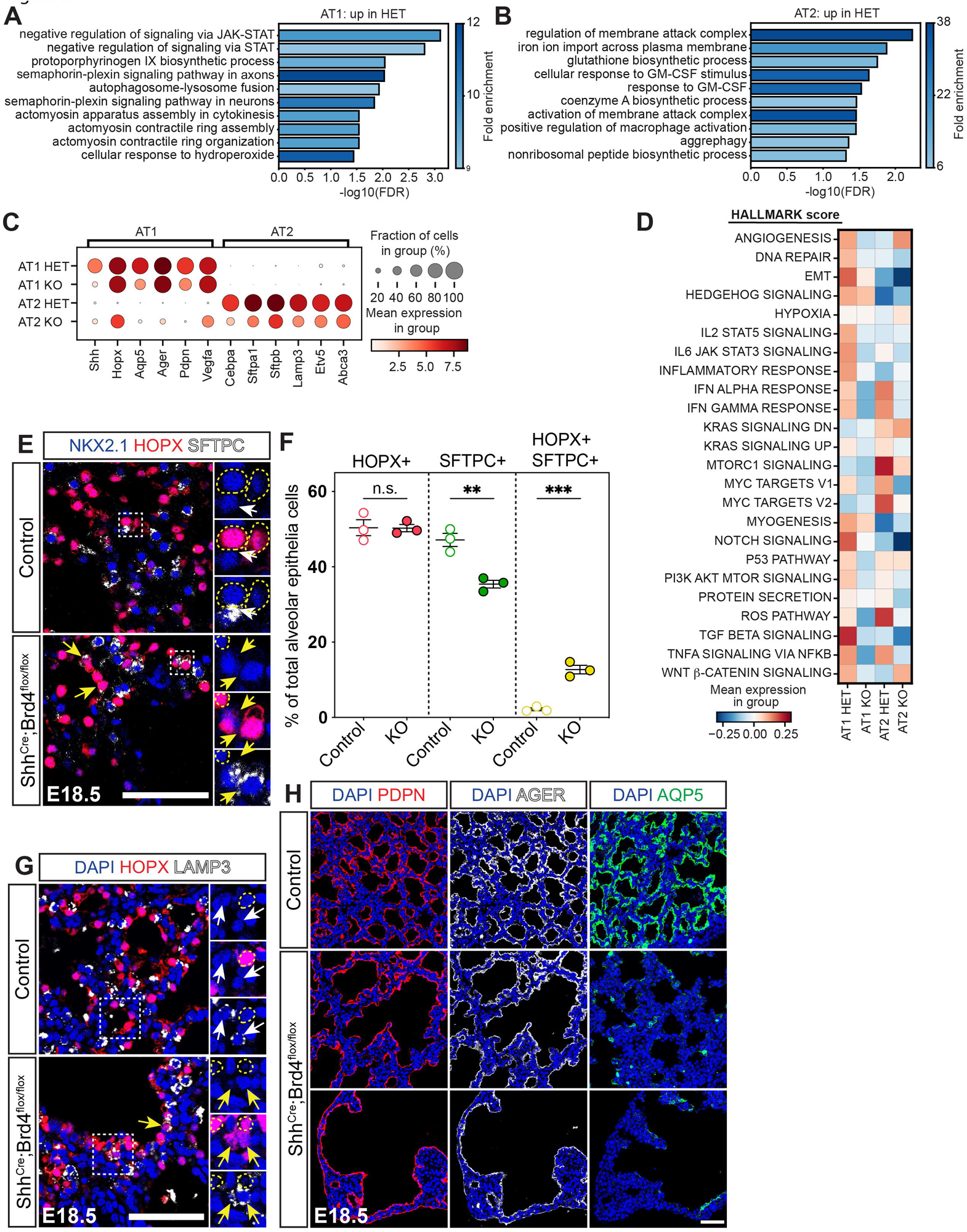
BRD4 is critical for AT2 lineage determination and AT1 cell maturation. (A-B) GO analysis for top 10 ranked categories using upregulated (UP) gene enrichment sets in cell populations. (A) GO analysis for AT1 epithelial cells, (B) GO analysis for AT2 epithelial cells. (C) Dot plots of expression of AT1 and AT2 cell marker genes. Dot size indicates the proportion of cells within a cluster expressing a gene, and color intensity indicates the relative expression level. (D) Heatmap representing expression levels of signaling and response pathway hallmark scores. (E) IHC for NKX2.1, HOPX, and SFTPC in control and mutant E18.5 lungs. Dashed yellow circles mark AT1 cells, white arrows mark AT2 cells, and yellow arrows mark HOPX+ SFTPC+ cells. (F) Quantification of AT1, AT2, and HOPX+ SFTPC+ cells at E18.5 in control versus mutant lungs as shown in panel (E). Quantification data are represented as mean ± SEM. Two-tailed t tests: n.s.: not significant, **p ≤ 0.01, ****p ≤ 0.001, n = 3. (G) IHC for HOPX and LAMP3 in control and mutant E18.5 lungs. Dashed yellow circles mark AT1 cells, white arrows mark AT2 cells, and yellow arrows mark HOPX+ LAMP3+ cells. (H) IHC for PDPN, AGER, and AQP5 in control and mutant E18.5 lungs. All scale bars: 50μm unless otherwise indicated. Dashed white boxes mark zoomed in areas.

To confirm our bioinformatic findings of impaired alveolar epithelial cell differentiation, we performed histological analysis on control and BRD4 mutant lungs at E18.5. Consistent with loss of AT2 cell capture in our scRNA-seq data set, we observed a loss of AT2 cells as a percentage of total alveolar epithelial cells (**Figures 6E-F**). Moreover, we observed a loss of distinct AT2 cell fate as a subset of cells expresses both SFTPC and HOPX, suggesting poor AT2 cell lineage determination (**Figures 6E-F**). These cells in the alveolar compartment did not express the more mature AT2 cell marker LAMP3, but rare exceptions of LAMP3+ HOPX+ cells were present in cystic distal airway structures (**Figure 6G**). Similarly, BRD4 deficient AT1 cells failed to mature. While they expressed PDPN and AGER in the alveolar compartment and within the cystic distal airway structures, BRD4 deficient AT1 cells exhibited minimal AQP5 expression (**Figure 6H**). However, despite the absence of mature AT1s that communicate with the capillary bed, capillary endothelial development appeared mostly unaffected in BRD4 mutant lungs (**Figures S6C-F**). Together, our data show that BRD4 is required for alveolar epithelial lineage determination and maturation.

## DISCUSSION

In this study, we determined that BRD4 is a critical regulator of lung development. Loss of BRD4 results in perinatal lethality accompanied by abnormal branching morphogenesis, proximal-distal patterning, and cell differentiation and maturation within the lung. This is largely due to disruption of the normal epithelial-mesenchymal crosstalk required for patterning and branching morphogenesis. This is some of the first evidence for epigenetic regulation of tissue crosstalk governing early lung development.

### Epithelial-mesenchymal crosstalk

Multiple loss-of-function genetic studies have identified cystic distal lung structures as a major phenotype.^6, 8, 9, 14, 34, 35, 40–43^ However, no clear, comprehensive mechanism driving this phenotype exists. A SHH-BMP4-FGF signaling loop plays an important role in branching morphogenesis.^44^ SHH expression in the distal tip endoderm increases proliferation and induces expression of BMP4 in adjacent, central mesoderm. BMP4 inhibits growth of the central, distal tip but allows for proliferation and extension of the lateral portions of the lung bud towards sources of FGF10 to form new buds.^36, 37^ Interrogation of this pathway in most studies with cystic distal lung structures revealed no changes in the endodermal-mesodermal patterning by SHH, BMP4, and FGF10. While loss of DICER, a global translational regulator, in lung endoderm leads to cysts and expansion of BMP4 and FGF10 in the mesoderm, the mechanism within endoderm governing epithelial-mesenchymal interactions is unclear.

BRD4 deficiency in the endoderm results in a reduction of distal tip SHH expression concomitant with an increase and expansion of BMP4 and FGF10. Loss of BRD4 disrupts normal BMP4 and FGF10 expression patterns, preventing sufficient formation of new lung buds. Similarly, SHH deficient embryos have expression of BMP4 and FGF10 at the distal lung buds, indicating a BRD4-SHH genetic interaction governing branching morphogenesis. Accordingly, SHH treatment of BRD4 deficient embryonic lung explants partially rescues the phenotype by reducing distal cyst size and expanding SOX2 expression. These findings suggest a BRD4-SHH interaction not only mediates epithelial-mesenchymal communication but also ensures appropriate expansion of proximal, SOX2+ endoderm during morphogenesis. Whether this BRD4-SHH interaction is direct is unclear as mesenchymal WNT5A deficiency also results in decreased SHH and expanded BMP4 and FGF10.^41^ As such, interrogation of the WNT pathway may provide additional details to the mechanism behind BRD4-mediated branching and proximal-distal patterning.

### Cell specification, identity, and maturation

Beyond regulation of morphogenesis, BRD4 is an important regulator of cell specification and differentiation during lung development. Besides its implication in heart disease and cancer, BRD4 plays a broad role in embryogenesis.^45–47^ BRD4 orchestrates implantation and embryonic growth, maintains stem cell identity, regulates fat and muscle development, and promotes neural crest and other progenitor cell differentiation.^23, 25, 26, 32^ We observed a cell autonomous defect in specification, differentiation, and maturation of both proximal airway and distal alveolar epithelial cell subtypes, indicating a global role for BRD4 in lung cell specification. Interestingly, loss of secretory cells in the conducting airway is more pronounced. Similar deficits in secretory cells after loss of BRD4 have been observed in the small intestine, suggesting a conserved mechanism for secretory cells specification.^48^ Whether BRD4 is a direct regulator of the genes important for specification of every type of lung epithelial is unclear. It is possible that early defects in proximal-distal patterning and branching disrupt alveolar differentiation in development.^49^ Future studies aimed at uncovering BRD4 genome occupancy during lung development may provide some clarity.

Cell non-autonomous differentiation defects are apparent as well with ectopic smooth muscle surrounding the entire distal cystic structure. We hypothesize that there is a breakdown in epithelial-mesenchymal crosstalk. Previously, both FGF and WNT signaling have been implicated in this process.^3, 50^ As such, it will be important to determine whether BRD4 could be regulating this process in a cell autonomous and/or non-autonomous role.

### Conclusion

These studies provide evidence of the role of epigenetic factor, BRD4, in lung development. Like previous studies on the epigenetics of lung development, there are global defects including deficits in branching morphogenesis, proximal-distal patterning, and progenitor cell specification and cell maturation.^5, 9, 11, 14^ Whether BRD4 plays a role in the processes of injury and repair across the lifespan is unknown. Current studies indicate BRD4 as a prominent participant in progression of cancer, heart disease, and lung fibrosis. Our data demonstrating BRD4 is a master regulator of lung development are important given ongoing clinical trials for cancer and fibrosis. Fetal programs are often reactivated for repair processes. In the setting of BRD4 inhibition, loss of its function could potentially be detrimental to repair after injury. Thus, future studies exposing its role in adult injury and repair are important.

### Limitations of the Study

The experiments here used in vivo and ex vivo lung explant modeling to assess the role of BRD4 in lung development. Loss of BRD4 disrupts epithelial-mesenchymal crosstalk that we believe is important for all aspects of development. These studies are limited in fully characterizing the pathways downstream of BRD4 regulation in that we are unable to perturb both epithelium and mesenchyme simultaneously. As such, we used lung explant modeling to reverse the BRD4 phenotype and were able to partially rescue the phenotype. There are likely other signaling pathways such as WNT working in conjunction to mediate this crosstalk. An additional limitation to these studies is defining the specific molecular mechanism of BRD4-dependent regulation of gene expression for epithelial-mesenchymal crosstalk. BRD4 is a multifunctional tool possessing the ability to regulate a broad category of genes. Regulation of gene transcription is mediated through both genetic and epigenetic mechanisms. These include the ability to read histone lysine residues, facilitate genome folding, occupy enhancers to recruit transcription factors, and directly activate transcription as a regulator of the P-TEFb/RNA polymerase II complex. Thus, a focus on the global function of BRD4 is more prudent and should guide further studies on the role of BRD4 across the lifespan in lung development, disease, and regeneration.

## ACKNOWLEDGEMENTS

We are grateful for the Ayla Gunner Prushansky Research Fund for support of this study. We thank Jarod Zepp for critical discussion and Keiko Ozato and Anup Dey for providing the *Brd4^flox^* mice. We thank the CHOP Flow Cytometry core led by Dr. Florin Tuloc for expertise and assistance to complete this study. This work was supported though the Parker B. Francis Foundation (D.B.F), Burroughs Wellcome Foundation Career Award for Medical Scientists (R.J.), and grants from the National Institutes of Health: K08 HL140129 (D.B.F.), R35 HL166663 (R.J.).

## AUTHOR CONTRIBUTIONS

Conceptualization, D.B.F.; Methodology, D.C.L., H.W., and P.C.; Software, N.S.M.; Validation, D.C.L., H.W., K.Q., and J.P.; Formal Analysis, D.C.L., H.W., P.C., and N.S.M.; Investigation, D.C.L., H.W., P.C., K.Q., J.P., A.J., and M.L.; Resources, R.J., and L.R.Y.; Data Curation, D.C.L., H.W., P.C., K.Q., J.P., and N.S.M.; Writing-Original Draft, D.C.L., H.W., and D.B.F.; Writing-Review & Editing, D.C.L., H.W., P.C., K.Q., J.P., N.S.M., A.J., M.L., M.P.D.N., R.J., and D.B.F.; Visualization, D.C.L., P.C., and H.W.; Supervision, D.C.L., H.W., and D.B.F.; Project Administration, D.C.L., H.W., and D.B.F.; Funding Acquisition, R.J., L.R.Y., and D.B.F..

## DECLARATION OF INTERESTS

All authors declare no competing interests.

**Figure S1: Perinatal lethality in BRD4 mutant mice**

(A) Table showing perinatal survival from 6 litters of a *Shh^Cre^;Brd4^flox/+^* male crossed with *Brd4^flox/flox^* females.

(B) Control and mutant pup images just after birth.

(C) Lung float test for control and mutant mice.

**Figure S2: Cellular changes associated with BRD4 mutant mice**

(A) IHC for NKX2.1 and cleaved caspase 3 (cCASP3) in control and mutant E18.5 lungs. Scale bars: 50μm.

(B) RNA FISH of *Etv5* and *Fgfr2* in control and mutant E18.5 lungs. Scale bars: 50μm.

**Figure S3: Fgfr2 inhibitor does not rescue BRD4 mutant cystic distal airway structure phenotype**

(A) *Ex vivo* lung explant brightfield images of control and mutant lungs at E12.5 (left) and E15.5 (right) with treatments: DMSO and Alofanib (ALOF) (scale bar: 500μm).

(B) Left: wholemount immunostaining for SOX2 and SOX9 after 3 days of ex vivo culture (scale bar: 500μm). Dashed white boxes indicate branching tips in control and mutant lungs. Right: zoomed in images of branching tips, dashed white lines outline a distal airway (scale bar: 250μm).

**Figure S4: Ectopic basal cells in BRD4 mutants**

(A) IHC for the basal cell marker, P63. Scale bars: 100μm.

**Figure S5: Alterations in gene expression in BRD4 mutant secretory and multiciliated cells**

(A) Cell compositional changes in BRD4 heterozygous and mutant lungs based on cell number.

(B) Dot plot of marker genes for multiciliated and secretory cell type where dot size indicates the proportion of cells within a cluster expressing a gene, and color intensity indicates the relative expression level.

(C-F) GO analysis for top 10 ranked categories using upregulated (UP) gene enrichment sets in cell populations. (C) GO analysis for secretory cells in heterozygous lung, (D) GO analysis for secretory cells in the mutant lung, (E) GO analysis for ciliated cells in the heterozygous lung, and (F) GO analysis for ciliated cells in mutant lung.

(G) Heatmap representing expression levels of signaling and response pathway hallmark scores.

**Figure S6: BRD4 mutant lungs maintain vascular density**

(A-B) GO analysis for top 10 ranked categories using upregulated (UP) gene enrichment sets in cell populations. (A) GO analysis for AT1 epithelial population in the mutant lung, (B) GO analysis for AT2 epithelial population in the mutant lung.

(C) IHC for EMCN (left) and ERG and CAR4 (right) in control and mutant E18.5 lungs. Scale bars: 50μm

(D-F) Quantification of capillary density and endothelial cell number in control and mutant E18.5 lungs as shown in panel (C). Data are represented as mean ± SEM. Two-tailed t tests: n.s.: not significant, *p ≤ 0.05, n ≥ 3 for each group.

## STAR METHODS

### RESOURCE AVAILABILITY

#### Lead contact

Requests for further information and resources should be directed to the lead contact, David Frank.

#### Materials availability

This study did not generate new unique materials.

#### Data and software availability

Raw and processed next-generation sequencing data sets have been uploaded to the NCBI GEO database (https://www.ncbinlm.nih.gov/geo/) and will be made public upon peer-reviewed publication.

### EXPERIMENTAL MODEL AND SUBJECT DETAILS

#### Mouse lines

Information related to the generation and genotyping of the following mouse lines has been previously described: *Shh^Cre^* (Jackson Laboratory stock # 005622)*, Brd4^flox^, and R26R^EYFP^* (Jackson Laboratory stock # 007903). The *Brd4^flox^* mice were generously provided by Keiko Ozato’s laboratory.^32^ Mice were maintained on a mixed C57BL/6 and CD1 background. Experiments were performed with a minimum of three animals per condition of mixed sex, and littermate controls were included. Controls included *Shh^Cre^* positive and negative animals. Animal procedures were ethically performed and approved under the guidance of the University of Pennsylvania and Children’s Hospital of Philadelphia Institutional Animal Care and Use Committee.

### METHOD DETAILS

#### Tissue processing and histology

Embryonic lungs were harvested in cold PBS, placed into 2% paraformaldehyde, and fixed for either 5-6 hours (wholemount) or overnight (IHC) at 4°C. After fixation, lungs were washed with PBS at least four times, dehydrated in a series of ethanol washes (30%, 50%, 70%, 95%, and 100%), embedded in paraffin wax, and sectioned at a thickness of 6μm. Hematoxylin and eosin (H&E) staining was performed on slides with maximum surface area to observe for gross morphological changes.

#### Immunohistochemistry (IHC) and RNA fluorescence in situ hybridization (FISH)

Immunohistochemical staining was performed as previously described.^15, 51^ Briefly, after deparaffinization, slides were incubated with 3% H_2_O_2_ for 15 min to quench the endogenous peroxidases. The slides were then blocked using 5% donkey serum and incubated with primary antibody at 4°C overnight in predetermined concentrations (Table 1.). The presence of relevant proteins was visualized using Alexa fluor secondary antibodies (Table 2.). After secondary antibody staining, slides were mounted in Vectashield Antifade Mounting Medium (H-1000, Vector Laboratories) and images using Leica DMi8 confocal microscope. RNA FISH was performed using the RNAscope Multiplex Fluorescent V2 Assay (323100, Advanced Cell Diagnostics (ACDBio)) according to the manufacturer’s instructions. RNA 3-plex negative control probe (DapB) and mouse specific 3-plex control probe (Polr2a) were used for control sections. The RNA target probes included *Shh, Bmp4, Fgf10, Etv5 and Fgfr2* and mRNA transcripts were visualized using OPAL fluorophore reagents (Opal 540 FP1494001, Opal 570 FP11488001 or Opal 650 FP1496001, Akoya Biosciences). Images were acquired using a Leica DMi8 confocal microscope after slides were treated with DAPI and mounted with Prolong Gold Antifade Mountant (P36930, Thermo Fisher Scientific).

**Table 1.** List of primary antibodies

| S.NO. | Protein | Host | Catalogue | Company | Dilution |
| --- | --- | --- | --- | --- | --- |
| 1 | BRD4 | Rabbit | A301-985A100 | Bethyl Laboratories | 1:4000 |
| 2 | NKX2.1<br>(TTF1) | Rabbit | ab76013 | Abcam | 1:50 |
| 3 | NKX2.1<br>(TTF1) | Mouse | MS-699-P1 | Thermo Fisher Scientific | 1:100 |
| 4 | SOX2 | Goat | AF2018 | R&D Systems | 1:50 |
| 5 | SOX9 | Rabbit | Ab185966 | Abcam | 1:200 |
| 6 | Cleaved<br>Caspase 3 | Rabbit | MAB835 | R&D Systems | 1:100 |
| 7 | TUBB4 | Mouse | MU178-UC | BioGenex | 1:10 |
| 8 | HOPX | Mouse | sc-398703 | Santa Cruz | 1:100 |
| 9 | SFTPC | Goat | sc-7706 | Santa Cruz | 1:100 |
| 10 | SCGB1A1 | Rabbit | MA5-29780 | Thermo Fisher Scientific | 1:100 |
| 11 | SCGB1A1 | Rat | MAB4218 | NovusBio | 1:40 |
| 12 | TAGLN | Goat | Ab10135 | Abcam | 1:100 |
| 13 | ACTA2 | Mouse | A5228 | Millipore Sigma | 1:100 |
| 14 | LAMP3 | Rat | DDX0191P-100 | NovusBio | 1:100 |
| 15 | PDPN | Mouse | Ab10288 | Abcam | 1:200 |
| 16 | AGER<br>(RAGE) | Rat | MAB1179 | R&D Systems | 1:50 |
| 17 | AQP5 | Rabbit | Ab92320 | Abcam | 1:100 |
| 18 | ERG | Mouse | Ab214341 | Abcam | 1:100 |
| 19 | EMCN | Goat | AF4666 | R&D Systems | 1:200 |
| 20 | EMCN | Rat | 14-5851-82 | Invitrogen | 1:100 |
| 21 | CAR4 | Goat | AF2414 | R&D Systems | 1:100 |
| 22 | CDH1 (E-Cadherin) | Rabbit | 3195 | Cell Signaling Technologies | 1:100 |
| 23 | KRT5 | Rabbit | Ab52635 | Abcam | 1:100 |
| 24 | TP63 | Mouse | sc-8432 | Santa Cruz | 1:25 |

**Table 2.** List of secondary antibodies

| S.NO. | Secondary | Host | Catalogue | Company | Dilution |
| --- | --- | --- | --- | --- | --- |
| 1 | EdU Alexa Fluor 647 | NA | C10635 | Invitrogen | 1:100 |
| 2 | Anti-Rabbit IgG-488 | Donkey | A-21206 | Invitrogen | 1:250 |
| 3 | Anti-Mouse IgG-555 | Donkey | A-31570 | Invitrogen | 1:250 |
| 4 | Anti-Goat IgG-647 | Donkey | A-21447 | Invitrogen | 1:250 |
| 5 | Anti-Goat IgG-488 | Donkey | A-11055 | Invitrogen | 1:250 |
| 6 | Anti-Guinea Pig IgG-647 | Donkey | 706-605-148 | Jackson Immunoresearch Lab | 1:250 |

#### Wholemount immunofluorescence

Wholemount samples were prepared and stained as previously described.^51, 52^ E12.5 or E15.5 lungs were fixed in 2% paraformaldehyde for a minimum of five hours to overnight at 4°C. Following fixation, lungs were washed at least four times in PBS and stained with Sox2 (goat, R&D, AF2018, 1:50) and Sox9 (rabbit, Abcam, Ab185966, 1:200) in 5% donkey serum and 0.5% Triton X-100 in PBS (0.5% PBST) for 2-4 hours at room temperature then overnight at 4°C with gentle rotation. The next day, lungs were washed four times in 0.5% PBST at room temperature for 30 minutes each and then washed overnight in 0.5% PBST at 4°C with gentle rotation. The following day, the lungs were stained with secondary antibodies (donkey anti goat, and donkey anti rabbit, 1:250) in 5% donkey serum and 0.5% PBST and incubated at room temperature for two hours and then overnight at 4°C with gentle rotation. Images were acquired using a Leica DMi8 Thunder widefield microscope or a Leica DMi8 Confocal microscope.

#### Ex vivo lung explant assay

Lungs were harvested and cultured for 72 hours as previously described.^9, 53, 54^ Briefly, E12.5 lungs were dissected into cold PBS, washed at least twice in DPBS supplemented with Antibiotic-Antimycotic and once in DMEM/F12 supplemented with Antibiotic-Antimycotic, and placed onto 1-μm pore transwell inserts in DMEM/F12 supplemented with Antibiotic-Antimycotic. Purmorphamine (3µM, Tocris Bioscience, 4551), and Alofanib (RPT835) (1µM, Selleck Chemicals, S8754) treatments were mixed with culture media, added to predetermined lung explants, and refreshed every 24 hours. Lungs were imaged using a Leica DMi8 Thunder widefield microscope at 0 hour, 24 hours, 48 hours, and 72 hours.

#### Single cell RNA sequencing (scRNA-seq)

Lungs were dissected and dissociated for scRNA-seq as previously described.^15, 51^ Live, EPCAM+ cells were sorted into FACS buffer (1% fetal bovine serum and 10mM EDTA diluted in 1X phosphate buffered saline) using the BD FACSJazz sorter (BD Systems) and were subsequently centrifuged, washed, and resuspended in FACS buffer. A subset of cells was stained with Trypan Blue (BioRad, 145-0013) and counted on a brightfield microscope using a hemocytometer to assess live cell concentration. Approximately 16,500 cells from pooled mice population of *Shh^Cre^;Brd4^flox/+^*heterozygous control or *Shh^Cre^;Brd4^flox/flox^* homozygous knockout mice were loaded into individual lanes of a 10X Genomics 3’ gene expression scRNA-seq assay (v3.1) chip (10X Genomics) to target recovery of 10,000 live cells. Captured mRNA was reverse-transcribed, amplified, and prepared for high-throughput sequencing on the Illumina HiSeq2500 platform according to manufacturer specifications, with a targeted read depth of 50,000 reads per cell.

### QUANTIFICATION AND STATISTICAL ANALYSIS

#### Alveolar area and diameter measurements

Morphometric analysis of alveolar area and diameter was performed on H&E images of alveolar areas acquired using a Leica Thunder widefield microscope. Alveolar areas and diameters were extracted with previously described MATLAB software.^55, 56^ For each mouse, the mean area and diameter were calculated from at least six randomly selected 20x fields.

#### Quantification of branching points

The branching defect analysis was performed on wholemount images of E12.5 lung buds obtained using a Leica thunder widefield microscope. Based on the Sox9 staining of the branches stemming from the two primary bronchi, we counted the number of branching points in FIJI.

#### Quantification of epithelial cell proliferation

To quantify proliferative epithelial cells, we adopted our previously established procedures using EdU staining.^51, 57^ Briefly, before the harvest of lung tissues, pregnant dams were injected intraperitoneally with 50 gm/kg of EdU, and the embryos were harvested 4 hours following the injection. After performing the click chemistry-based EdU proliferation assay (Click-iT Plus EdU Alexa Fluor 647, C10640, Thermo Fisher Scientific), images of airway regions were captured using a Leica DMi8 confocal microscope. The percentages of proliferative cells in proximal and distal airways were counted using cell counter plugin in FIJI based on SOX2+/EdU+ and SOX9+/EdU+ respectively.

#### Quantification of ex vivo lung explant rescue

The tip number, tip thickness, and area of Sox2 expression were quantified based on wholemount immunofluorescence staining of Sox2 and Sox9. The IHC images of the whole lung explants were captured using a Leica DMi8 Thunder widefield microscope. All the tips of the lung explant marked by Sox9 staining were manually counted across the entire z stack using the cell counter plugin in FIJI. Due to the oval shape of the tips, straight lines were drawn across the shorter minor axis of the tips to measure the tip thickness in FIJI. The thickness of over 50 lung tips was measured and averaged for each lung explant. For measure the area of Sox2 expression, images of Sox2 staining were turned into binary using the threshold automatically calculated within FIJI. Afterwards, areas of Sox2 expression were selected using the wand tool, added onto the ROI manager, and measured.

#### Quantification of capillary endothelial defect

The capillary defect was characterized using IHC staining for pan-endothelial marker ERG (mouse, ab214341, Abcam, 1:100), capillary and venous endothelial marker EMCN (rat, 14-5851-82, Invitrogen, 1:100), and CAR4 (goat, AF2414, R&D Systems, 1:100), which marks a novel population of highly proliferative endothelial cells.^58^ Images were then processed using Imaris software (version 10.0.0). 5 identical distal lung images per mouse were randomly captured using a Leica DMi8 confocal microscope. Macrovasculature was excluded in these images to focus only on capillaries. The relative loss of endothelial cells was calculated based on the ratio of ERG+ cells in relation to DAPI+ cells in the field. The ‘colocalization’ function in Imaris was used to construct a new colocalized channel based on the signals of ERG and DAPI staining. To automatically quantify the number of ERG+ DAPI+ cells and the total number of DAPI+ cells, the ‘cell’ function in Imaris was used based on the colocalized channel and the channel of DAPI respectively. Quantification of capillary density was performed based on previously determined procedures.^51^ Volume of the whole tissue (DAPI), capillary EC (EMCN), and Car4-high capillary EC (Car4) was measured using the ‘surface’ function and stored under volume statistics. The capillary density and Car4-high capillary density were calculated by dividing the capillary EC volume and Car4-high capillary EC volume by the total lung volume respectively.

#### Cell Counting

Cell counting analysis was performed based on previously established protocol.^57^ Briefly, IHC images were captured using a Leica DMi8 confocal microscope and processed in FIJI. The cells of interest were manually counted based on signal colocalization using the cell counter plugin in FIJI. For a given stain, at least 500 cells or five confocal stacks captured at 40x magnification selected images were counted per animal.

#### Single-cell RNA sequencing analysis

Reads were aligned to the GRCm38/mm10 mouse genome, and unique molecular identifiers (UMIs) were counted on a per-cell, per-gene basis using STAR-solo.^59^ Putative cells were separated from ambient background RNA using EmptyDrop, as implemented in STAR-solo.^59^ Downstream scRNA-seq processing, including filtering, normalization, cell type identification, cluster identification, and differential gene expression analysis, was performed using scanpy.^60^ Gene expression counts were normalized to CPM using the sc.pp.normalize_total function and log-transformed to ln(1+CPM) using the sc.pp.lop1p function. When used in this work, Z-scored gene expression was calculated by standard-scaling ln(1+CPM) expression values using the sc.pp.scale function.

Cells were filtered to only include those with greater than 1000 unique genes detected and fewer than the 95th percentile (4714 genes) of unique genes detected to remove low-efficiency cell capture droplets and cell capture droplets possibly containing multiple cells, respectively. Likewise, cells with more than the 95th percentile of percentage of UMIs derived from mitochondrial mRNAs (11.5%) were excluded due to the high likelihood of those cell capture droplets containing unhealthy or dying cells. Cell doublets were further identified and removed using the scVI-tools implementation of the SOLO doublet detection algorithm.^61, 62^

Batch-corrected principal component analysis (PCA) was performed using the pytorch implementation of the harmony algorithm.^63^ The resultant PCA representation of the combined dataset was used to generated a k-nearest-neighbors network of cells using the sc.pp.neighbors function, with n_neighbors set to a value of sqrt(0.25 * n_cells) (29 cells), which is closely related to the heuristic proposed in.^64^ A Uniform Manifold Approximation and Projection (UMAP) was calculated for plotting purposes using the sc.pp.umap function, with init_pos=”spectral”. Cell clustering was performed using the sc.pp.leiden function with resolution=1 unless otherwise specified.^65–67^ Cells were assigned to cell types based on the expression of canonical marker genes for lung AT1, AT2, secretory, and multiciliated cell types.

Differential expression testing was performed using the sc.tl.rank_genes_groups function with method=”wilcoxon”. Differentially expressed genes were filtered to include only those with a log2-fold change (LFC) of magnitude greater than 1 and were additionally filtered to remove those genes associated with sex-chromosome regulatory mechanisms, such as the Xist gene, as well as pseudogenes, mitochondrial genes, ribosomal genes, and other genes that are not of interest in this study.

Over-representation analysis (ORA) of differentially expressed genes was performed using gProfiler, with query genes being ordered by LFC in order of descending magnitude.^66^ Term fold-enrichment was calculated as the number of queried genes in the given term (intersection_size in gProfiler) divided by the number of genes expected to be in the given term by random chance (query_size * term_size / effective_domain_size in gProfiler). Over-represented terms with FDR<0.05 were first ordered by term fold-enrichment in descending order, and the terms with highest term fold-enrichment were subsequently taken and ordered by ascending FDR before plotting. MSigDB hallmark gene set scoring was performed across subsets of cells by first z-scoring the ln(1+CPM) gene expression values of those cells and then calling sc.tl.score_genes function from scanpy on genes contained in each gene set.^68, 69^

#### Statistics

Shapiro-Wilk normality test was first performed to determine whether the following statistical tests should be parametric or nonparametric. If the data passed the Shapiro-Wilk normality test, the parametric Student’s t tests were performed. Otherwise, the nonparametric Mann-Whitney U tests were conducted. The F tests were also used to determine whether equal variance should be assumed between groups. All previous statistical analyses were performed using the built-in analysis software in Prism 9.0 with the following statistical significance: ∗ = p ≤ 0.05, ∗∗ = p ≤ 0.01, ∗∗∗ = p ≤ 0.005, ∗∗∗∗ = p ≤ 0.001. Each point on the graph represents a single biological replicate obtained through averaging cell counts from multiple regions of a single animal.

