## Supplementary figures and images for "Endodermal BRD4 mediates epithelial-mesenchymal crosstalk during lung development"

### Figure S1

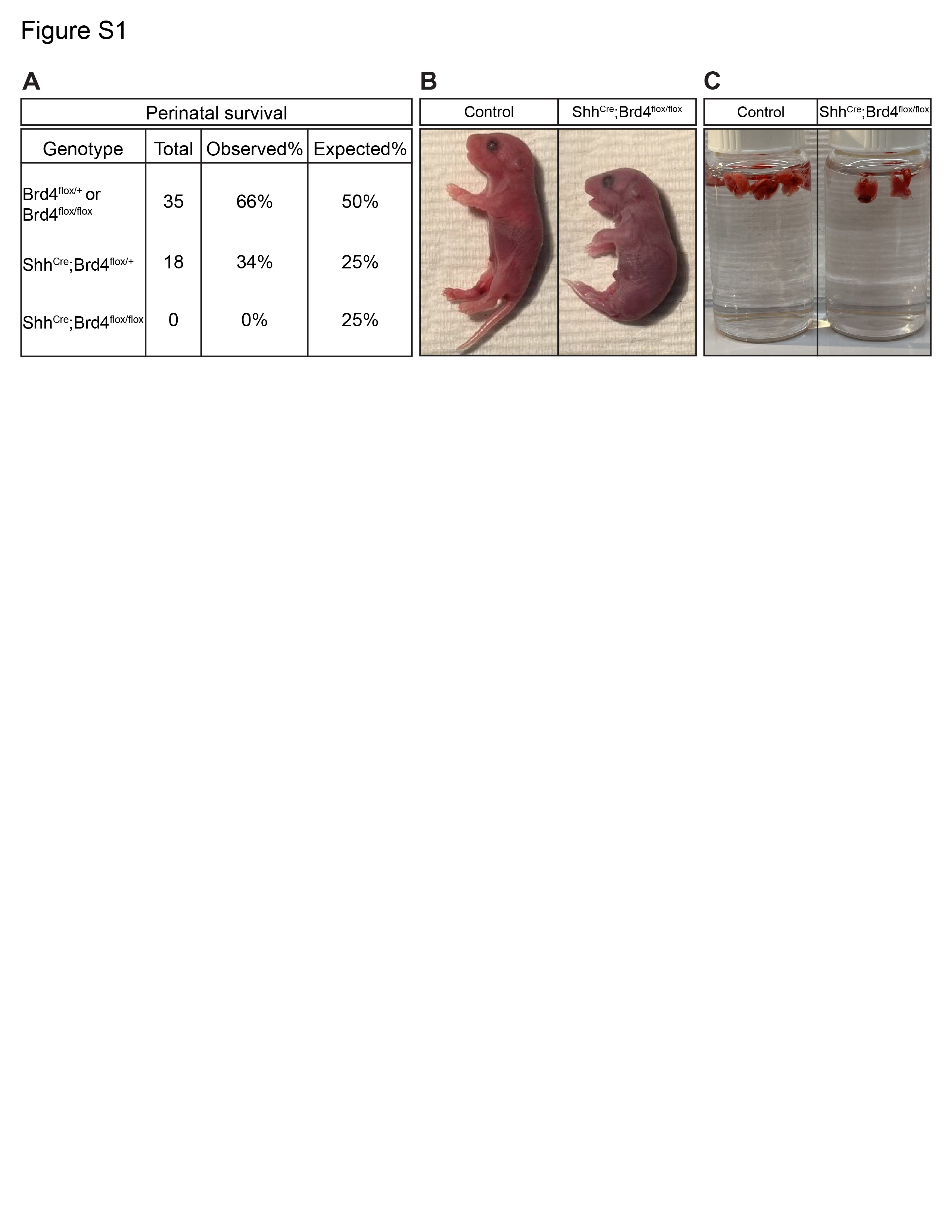

### Figure S2

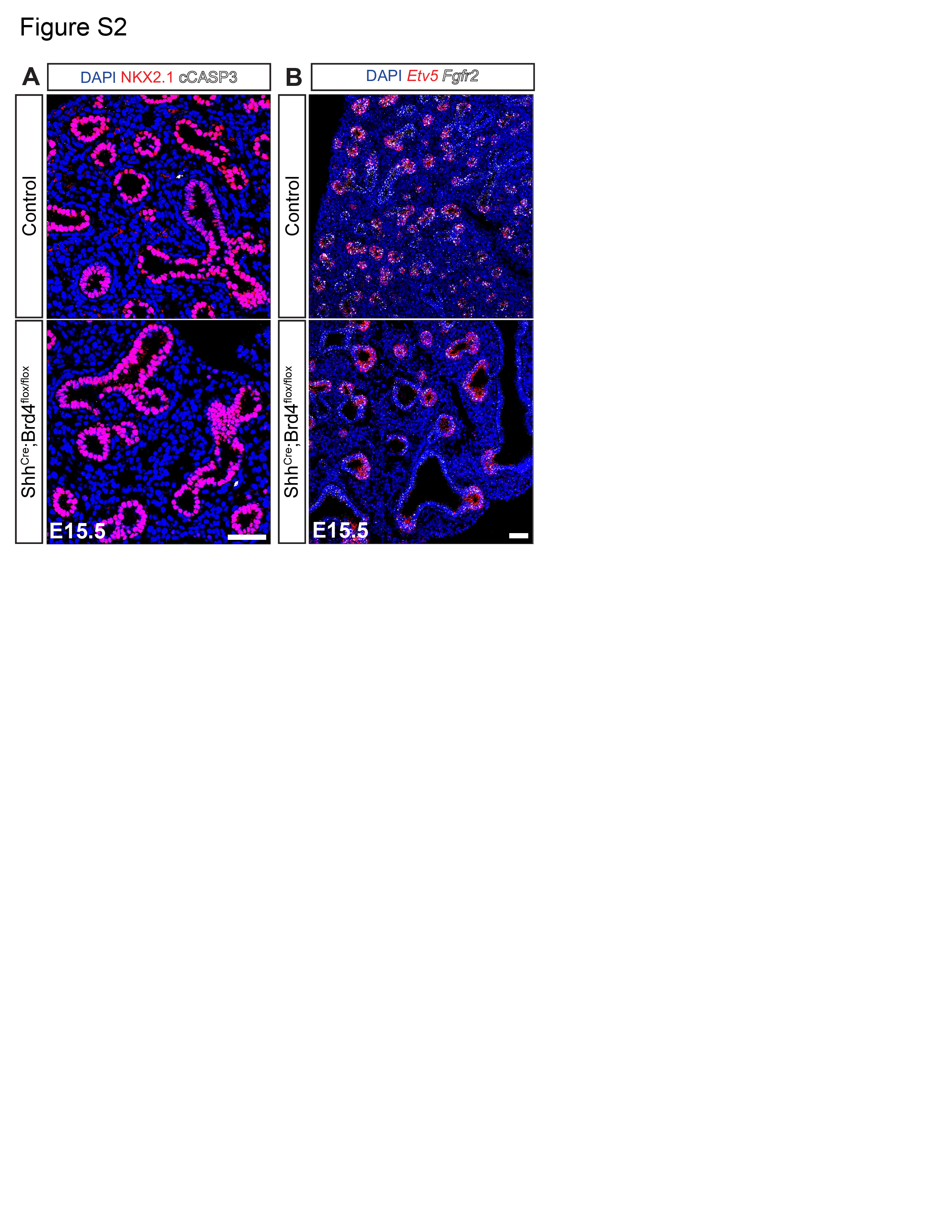

### Figure S3

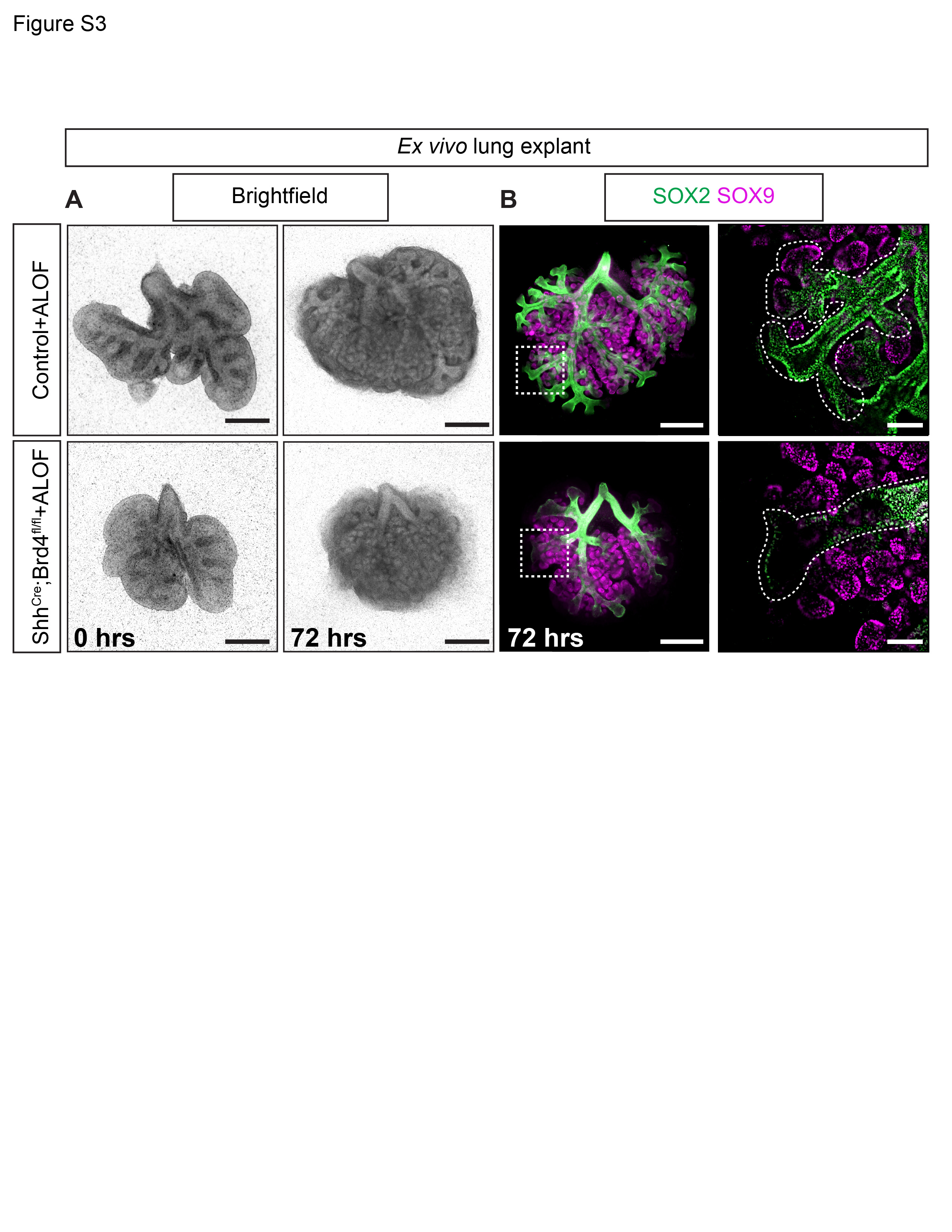

### Figure S4

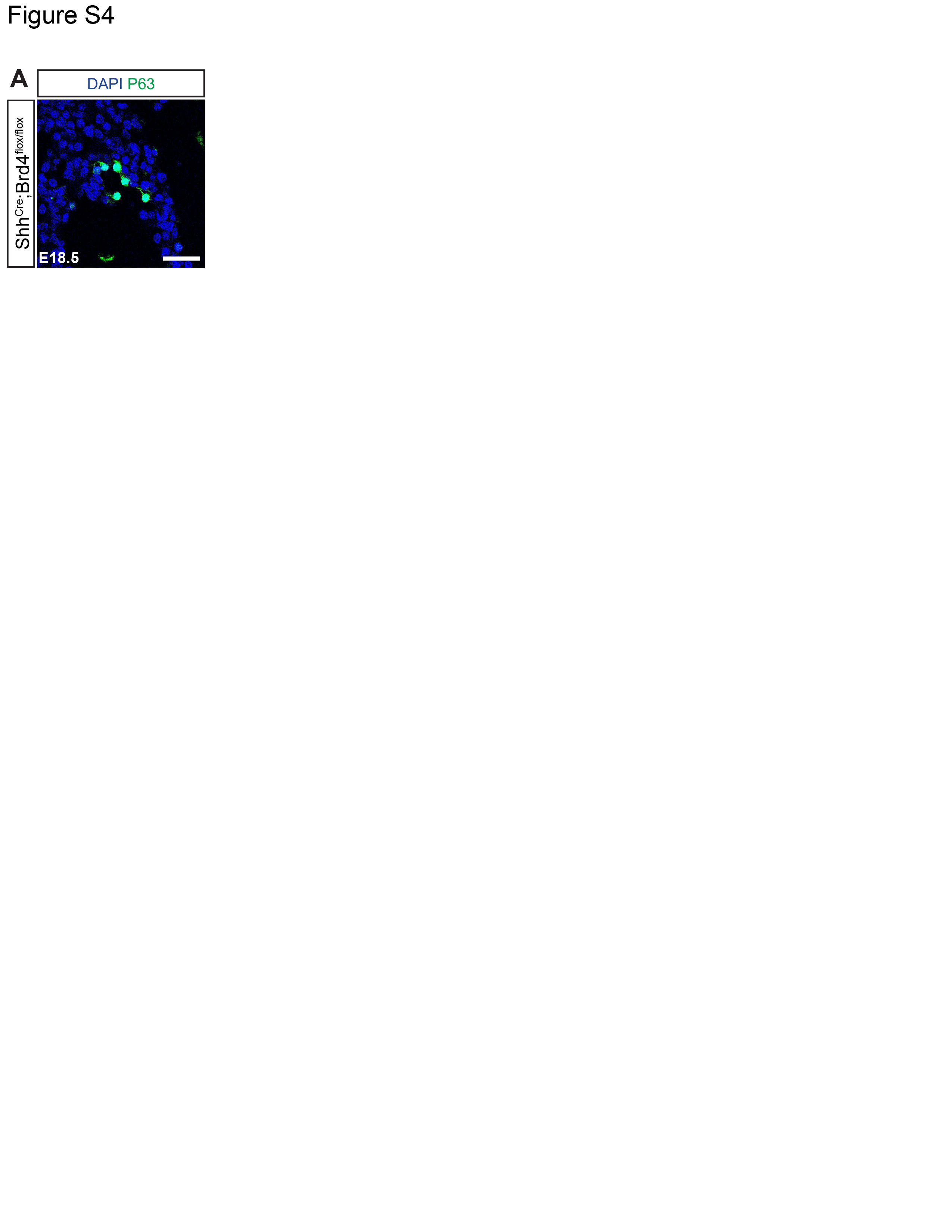

### Figure S5

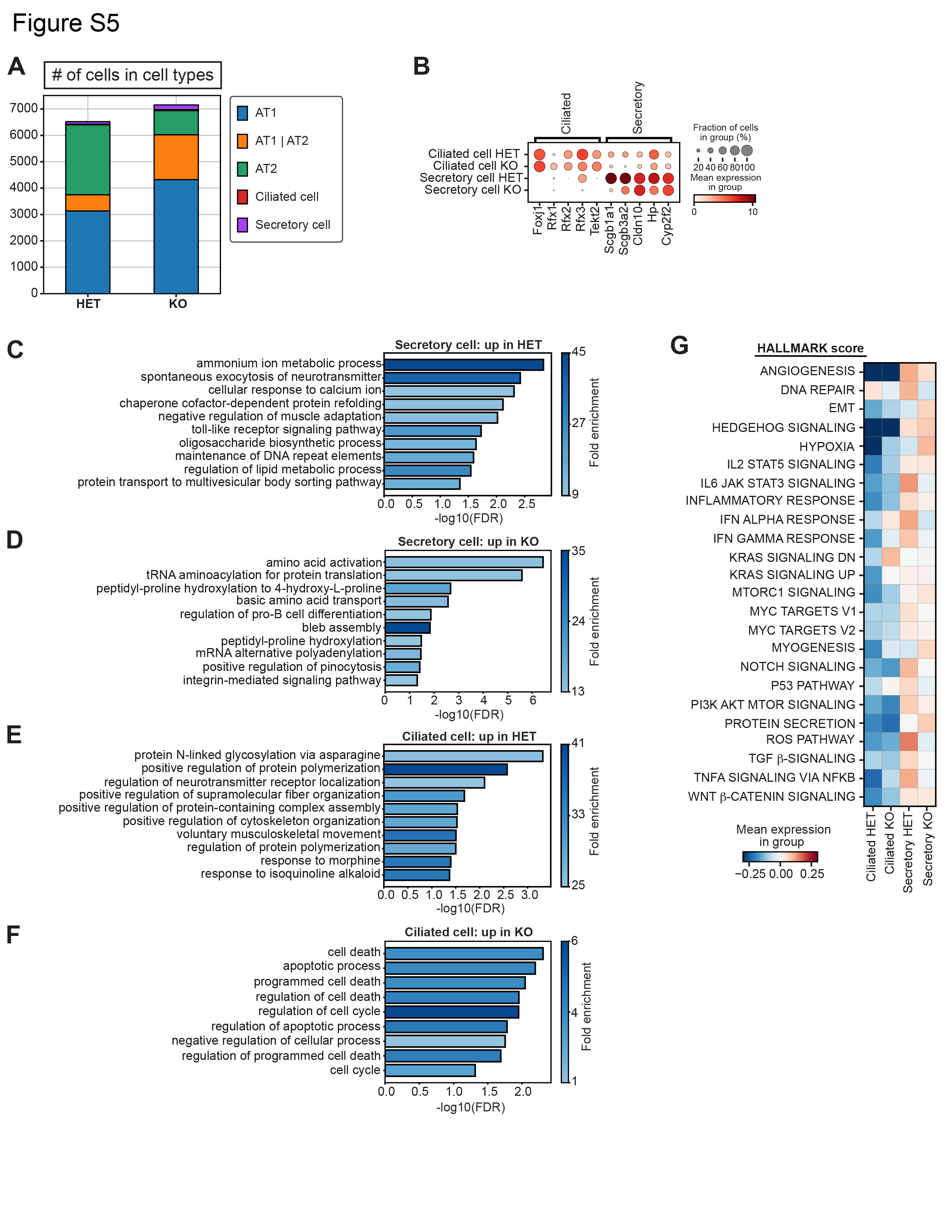

### Figure S6

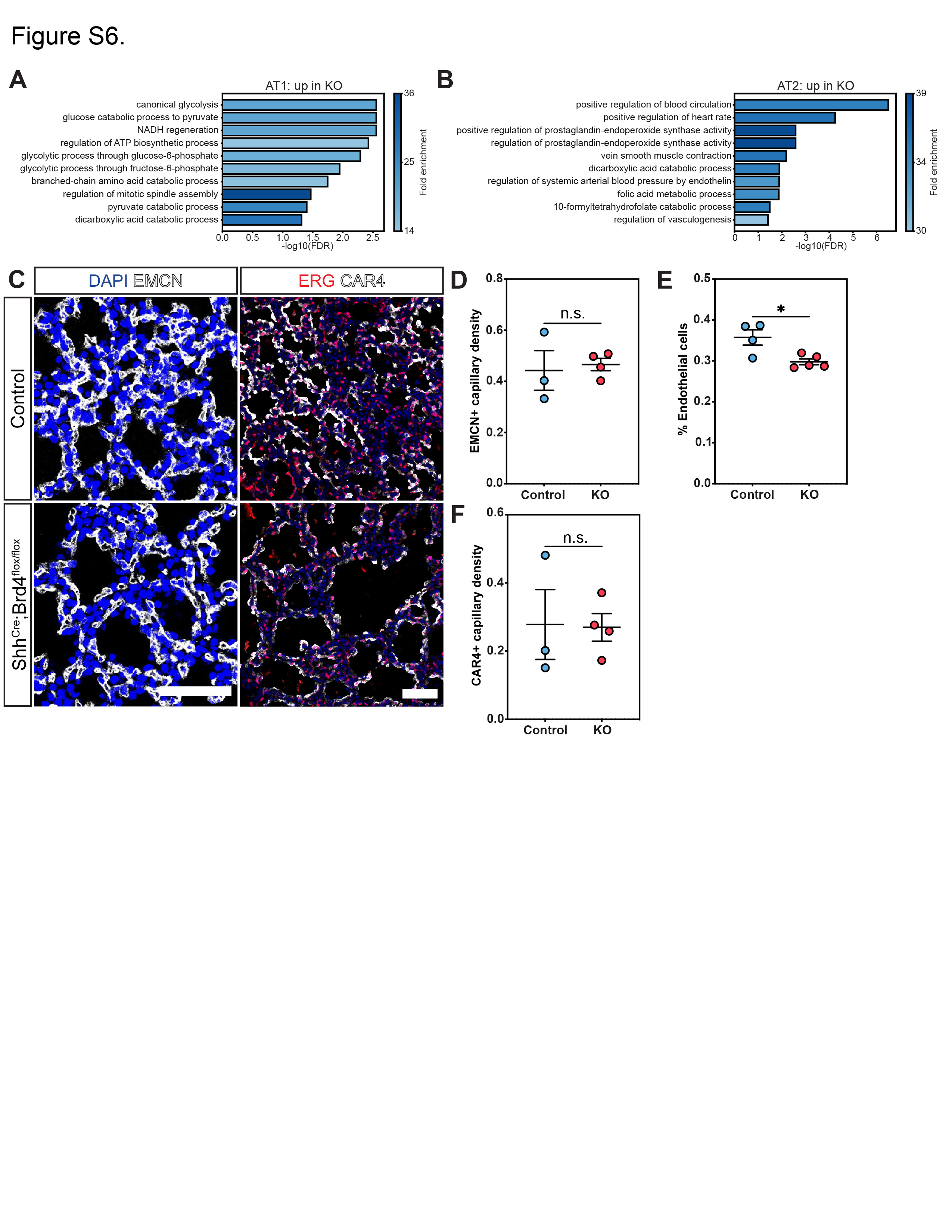
